# A globally threatened shark, *Carcharias taurus*, shows no population decline in South Africa

**DOI:** 10.1101/2020.06.02.130005

**Authors:** Juliana D. Klein, Aletta E. Bester-van der Merwe, Matthew L. Dicken, Arsalan Emami-Khoyi, Kolobe L. Mmonwa, Peter R. Teske

## Abstract

Knowledge about the demographic histories of natural populations helps to evaluate their conservation status, and potential impacts of natural and anthropogenic pressures. In particular, estimates of effective population size obtained through molecular data can provide useful information to guide management decisions for vulnerable populations. The spotted ragged-tooth shark *Carcharias taurus* (also known as the sandtiger or grey nurse shark) is widely distributed in warm-temperate and subtropical waters, but has suffered severe population declines across much of its range as a result of overexploitation. Here, we used multilocus genotype data to investigate the demographic history of the South African *C. taurus* population. Using approximate Bayesian computation and likelihood-based importance sampling, it was found that the population underwent a historical range expansion that may have been linked to climatic changes during the late Pleistocene. There was no evidence for a recent anthropogenic decline. Together with census data suggesting a stable population, these results support the idea that fishing pressure and other threats have so far not been detrimental to the local *C. Taurus* population. The results reported here indicate that South Africa could possibly harbour the last remaining, relatively pristine population of this widespread but vulnerable top predator.

## Introduction

The size of a population is a determining factor in how evolutionary and demographic processes affect its long-term survival. In conservation biology, small populations are of concern as they are typically more susceptible to demographic, environmental and stochastic genetic events^1–3^. The loss of genetic diversity and accumulation of deleterious alleles can lead to a decreased ability to adapt to changing environmental conditions, ultimately impairing long-term persistence and resulting in an elevated extinction risk^4,5^. The concept of the effective population size (*N*_e_) was originally introduced by Wright^6^ and can be interpreted as the size of an ideal population that experiences the same amount of genetic drift as the population under consideration. ‘Ideal’ in this context refers to a constant population of equal sex ratio, in which all individuals mate randomly, and are equally likely to produce offspring^6^. In wild populations, several factors, such as skewed sex ratio and non-random variance in reproductive success and mating, influence the amount of genetic diversity lost, and *N*_e_ is often reduced in comparison to actual, or census size (*N*_c_)^7–9^.

Several studies have attempted to establish the ratio of *N*_e_ and *N*_c_, with the expectation that the estimation of one parameter would allow the inference of the other^10^. Although this is an active area of research, ratios remain uncertain, as they are influenced by a variety of factors which interact in reducing *N*_e_. Reported average *N*_e_/*N*_c_ ratios across a range of species from different taxa range from approximately 0.1 to 0.5 ^11–13^. *N*_e_/*N*_c_ ratios in teleost fish with high fecundity, high juvenile mortality and sweepstake recruitment (type III survivorship), can be as low as 10^−5^ ^14,15^ (but see Waples *et al*.^16^). In contrast, in elasmobranchs (sharks, rays and skates), *N*_e_ closely approximates *N*_c_^17,18^. This is in line with theoretical predictions based on their life history traits such as longevity, late maturity and low fecundity rate^19^. Nevertheless, considerable variation exists among species, as shown by Chevolot et al.^20^, where *N*_e_/*N*_c_ ratios for the thornback ray *Raja clavata* were estimated to be very low (between 9 × 10^−5^ and 1.8 × 10^−3^).

Effective population size *N*_e_ can be estimated from genetic and demographic data, or a combination thereof^16,21,22^. As it is usually difficult to collect enough data on demographic parameters in wild populations, genetic approaches have become widely popular, and are particularly useful when studying rare or elusive organisms, such as elasmobranchs^23^. In light of current levels of global fisheries exploitation, estimates of contemporary *N*_e_ and demographic changes through time can provide valuable information to guide effective management and conservation of vulnerable elasmobranch species and populations^24–26^. As a coastal species, the ragged-tooth shark *Carcharias taurus* is not targeted by large off-shore fishing industries, but it is nonetheless threatened by small-scale fishing operations such as artisanal and recreational fisheries^27,28^. Population declines have been observed over much of the species’ distribution, and the species is listed as critically endangered in South America^29^ and East Australia^30^, and is thought to be locally extinct in the Mediterranean^31^. A large number of sharks has been caught in the beach meshing program along the Australian east coast, which is believed to be a significant driver in the dramatic decline experienced by *C. taurus* in that region^32,33^. In South Africa, the KwaZulu-Natal beach meshing program has also been identified as a having a negative impact on this species, due to its very low intrinsic rate of population increase^34^. Despite the potential impact of these threats, no decline in catch rate or body size of *C. taurus* was detected in the competitive shore-fishery^35^ or beach meshing catches^34^. Population trends estimated from mark-recapture data also remained stable over a 20-year study period^36^. In the same study, it was estimated that an average of 6,800 juvenile and 16,700 adult *C. taurus* inhabit the South African coast. However, this modelling approach was applied to a shark species for the first time, and some uncertainties remain around the parameters of the model^36^. To gain a better understanding of whether population trends inferred from catch data present an accurate picture of the health of this species, we used genetic data to explore the demographic history of the South African *C. taurus* population. In particular, we aimed to confirm that the population has not yet suffered any drastic declines from anthropogenic pressures. This represents the first genetic assessment of its kind for this population and may help to direct future management decisions.

## Methods

### Sampling and data generation

Genetic data for *Carcharias taurus* were generated using opportunistically collected samples originating from both juvenile and adult sharks. Research fishing trips were conducted at various nursery sites along the South African south coast from 2003 to 2009. Additional tissue samples from the east coast were provided by the KwaZulu-Natal Sharks Board beach meshing program. All capture and sampling protocols were conducted following guidelines endorsed by the Port Elizabeth Museums animal ethics committee. DNA extraction and amplification of 12 microsatellite markers was conducted following protocols described in Klein *et al.*^37^ for a total of 189 samples. MICROCHECKER 2.2.3^38^ was used to examine genotypes for stuttering, null alleles and large allele drop-out. Linkage disequilibrium (LD) between pairs of loci and significant deviations from Hardy-Weinberg-Equilibrium (HWE) were tested in GENEPOP 4^39^. For both tests, the dememorization number was set to 1,000, the number of batches to 500 and the number of iterations per batch to 10,000. P values were corrected for multiple comparisons with the B-Y method^40^ using the *p.adjust* function in R^41^. Because inferences of past population size changes can be strongly affected by genetic structure^42,43^, it was first confirmed that the species is genetically homogeneous in South African waters. For this purpose, population pairwise *F*_ST_ and *D*_est_ values were computed in GENALEX 6.5^44^. P values were generated under 1,000 permutations and adjusted using B-Y correction. Further, to visualize genetic distances among individuals, a principal coordinate analysis (PCoA) was conducted in GENALEX. Subsequent analyses were aimed at exploring the demographic history of *C. taurus* in South Africa using two complementary methods that evaluate temporal changes in *N*_e_.

### Approximate Bayesian computation

Approximate Bayesian computation (ABC) as implemented in DIYABC 2.0^45^ was used to compare several pre-defined models of historical change in *N*_e_. ABC is a Bayesian approach in which coalescent-based simulations are compared to observed data using summary statistics to identify models that fit the observed data most closely. Comparison of different demographic models was preceded by exploratory analyses to establish suitable priors. To this end, a scenario with a single, genetically homogeneous population was simulated (1 × 10^6^ data sets) to determine posterior distributions of *N*_e_ and mutation rate (μ). A full description of this analysis is provided in the Supplementary Information.

Following this preliminary analysis, three competing demographic scenarios were constructed: a stable population (scenario 1), a population expansion (scenario 2), and a decline in population size (scenario 3) (Figure 1). Priors for *N*_e_ and μ were refined based on posterior values resulting from the previous analysis. To explore the approximate time when changes in population size occurred, we performed two analyses using the same three models, but with different priors for the timing of population size change (*t*). First, a wide, uniform prior was chosen to encompass more recent events such as the arrival of European settlers and the start of commercial fisheries in South Africa, as well as natural historical events that preceded the onset of European settlement (analysis 1). To ensure that the genetic signal of a recent, anthropogenic population decline was not masked by more significant changes in population size earlier in the history of the population, the same analysis was repeated with the prior for *t* restricted to the onset of commercial fisheries until present (analysis 2). All prior values are listed in Supplementary Table S3. To convert time in generations to years, an average generation time of 9 –10 years was assumed^46^. A data set consisting of 3 × 10^6^ iterations was simulated (1 × 10^6^ simulations per scenario). Summary statistics calculated included mean number of alleles across loci, mean gene diversity across loci, mean allele size variance across loci, and the mean ratio between the number of alleles and the range in allele size (Garza-Williamson index^47^). To identify the most likely demographic scenario, relative posterior probabilities with 95% confidence intervals (CI) were estimated using a logarithmic method as described by Cornuet *et al.*^48^. A detailed description of the evaluation of model fit and reliability of model choice is provided in the Supplementary Information.

**Figure 1:**
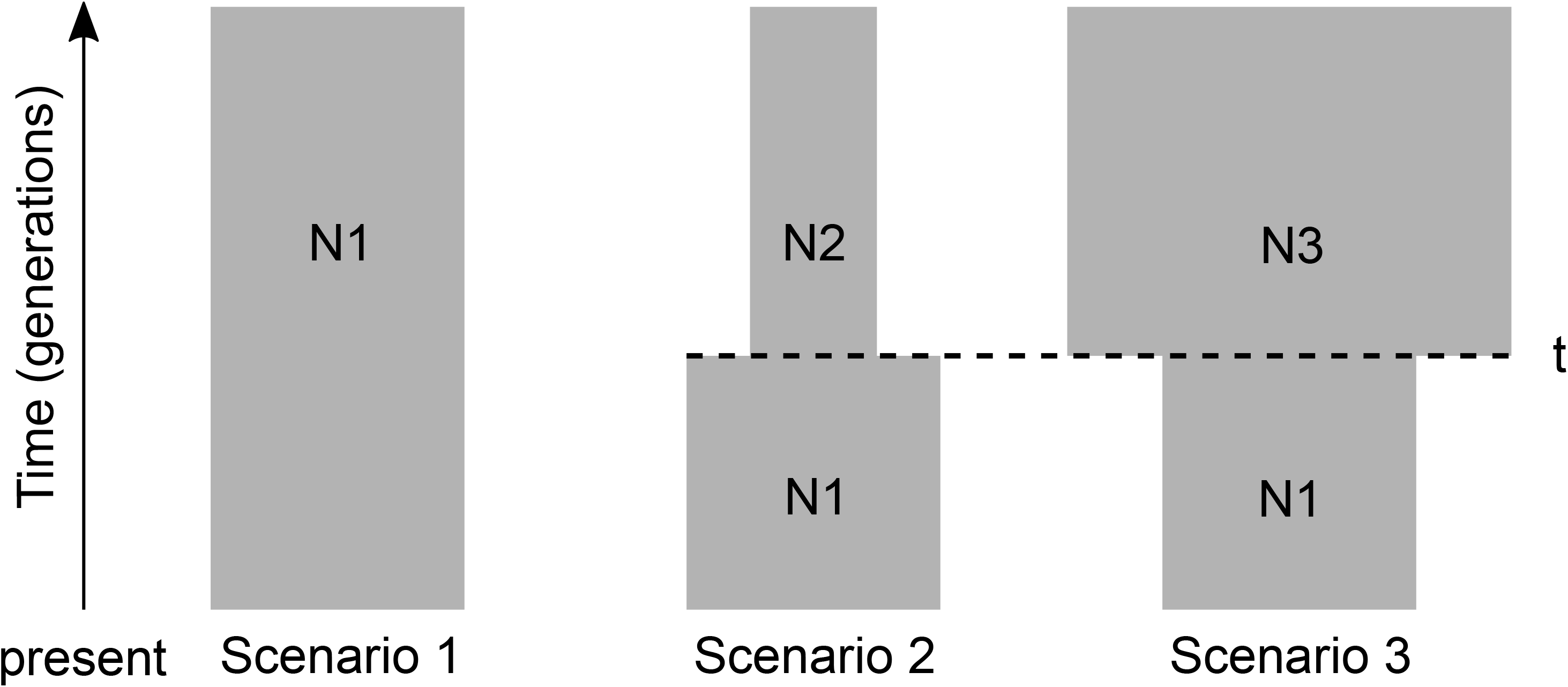
Demographic scenarios for *Carcharias taurus* tested using approximate Bayesian computation. N1 is the current effective population size and *t* is the time (in generations) at which the size of the population changed. N2 and N3 represent smaller and larger historical effective population sizes relative to N1, respectively. Prior distributions of each parameter are listed in Supplementary Table S3.

### Likelihood-based importance sampling

Likelihood-based importance sampling (IS)^43,49,50^ in MIGRAINE 0.5.4.1 (http://kimura.univ-montp2.fr/~rousset/Migraine.htm) was used as a complementary method to explore the demographic history of *C. taurus*. A combination of strict likelihood and PAC-likelihood (products of approximate conditionals^51^) computations (*PACanc* option) was used to infer parameters of interest under a model of a single population undergoing an exponential change in size, starting at some point in the past and continuing to the present. The generalized stepwise mutation model (GSM) was used with the parameter *P* describing the geometric distribution of stepwise mutations. Point estimates and corresponding 95% CIs were inferred for the following scaled parameters: current population size *θ* = 2*N*μ, ancestral population size *θ*_anc_ = 2*N*_anc_μ, where μ is the mutation rate per locus per generation and *N* is the number of gene copies per deme, and *D* = *Dg*/2*N*, an estimator of the time when population size started to change. The extra composite parameter *N*_ratio_ = *θ*/*θ*_anc_ was also estimated, which characterizes the strength of the population size change, with values <1 indicating a contraction and values >1 an expansion. Hence, significant departures from the expectations of a stable population were evident when 95% CI of *N*_ratio_ did not encompass 1^43^. To ensure that enough points were sampled near the top of the likelihood surface, 13 successive iterations were run with 500 parameter points and 200 simulated ancestral histories per point, followed by 8 iterations with 500 parameter points and 2,000 simulations per point. Scaled population size (*θ, θ*_anc_) and time (*D*) parameters were converted to effective population sizes (*N*_e_ and *N*_anc_) and time in years (*T*) using a microsatellite mutation rate of μ = 4.23 × 10^−4^ as estimated by DIYABC (see results), which is consistent with rates previously used in sharks^52,53^. As in the ABC analysis, a generation time of 9–10 years was assumed.

## Results

None of the microsatellite markers displayed stuttering, null alleles or large allele drop-out. No significant LD was detected between pairs of loci. One marker (*Ct243*) deviated significantly from HWE when all samples were analysed jointly (P value after BY correction: 0.026), but no deviations were observed when testing by sampling location. Population pairwise *F*_ST_ and *D*_est_ ranged from 0.005 to 0.020 and from 0 to 0.048, respectively (Supplementary Table S1) but none of the comparisons were significant after correction for multiple comparisons. The PCoA further indicated genetic homogeneity, with the first two axes only explaining 4.34% and 3.63% of the variation in the data, and overlap of groups from different sampling sites being evident (Supplementary Fig. S1). These results indicate that *Carcharias taurus* in South Africa comprises a single population and justify the inclusion of all available data in the subsequent demographic analyses.

### Approximate Bayesian computation

In the exploratory ABC analysis step, suitable prior ranges for the model comparison were successfully established (see Supplementary Information for results). For Analysis 1, which included a wide prior for the time at which the size of the population changed (*t*), scenario 2 (population expansion), had the highest posterior probability which was close to the maximum possible value of 1.0 with very narrow 95% CIs (Figure 2a). When *t* was restricted to the past ~400 years (analysis 2), Scenario 1 (constant population size) had the highest support when estimating posterior probabilities (Figure 2b). In both analyses, scenario 3 (population decline) received no support. Because simulations with *t* restricted to the last ~400 years did not provide conclusive posterior distributions as much of the genetic signal present in the data was excluded, estimates of demographic parameters were derived from scenario 2 simulated in analysis 1. Numerical values of posterior distributions with corresponding 95% highest posterior density intervals (95% HPDI) for all parameters estimated from scenario 2 are shown in Supplementary Table S4. A population size change occurred approximately 4,120 generations ago (95% HPDI: 669 – 9,420), corresponding to ~32,960 years, where *N*_e_ increased from 1,100 (95% HPDI: 134 – 5,810) to the current effective size of 15,900 (95% HPDI: 9,810 – 21,600). In addition, simulation-based evaluation demonstrated high power to distinguish between competing demographic models investigated here and validated this model as the most likely demographic scenario. A full description of these results is provided in the Supplementary Information.

**Figure 2:**
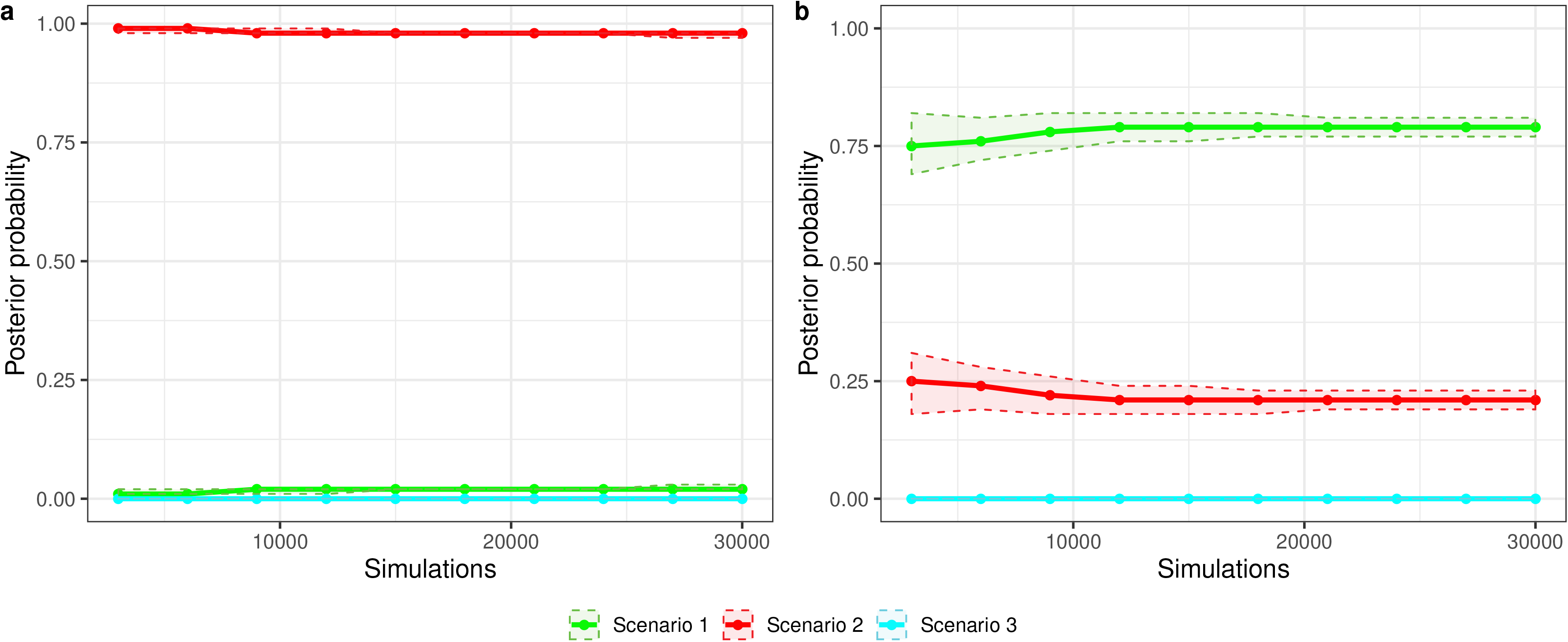
Estimation of relative posterior probabilities of the three scenarios based on a logistic regression-based estimate with (**a**) prior for *t* set to 1 – 5 × 10^3^ (analysis 1) and (**b**) prior for *t* restricted to 1 – 50 (analysis 2). Shaded areas indicate 95% confidence intervals. Y-axis is posterior probability, x-axis is number of simulated data sets closest to the observed data.

### Likelihood-based importance sampling

Inferred pairwise likelihood-ratio profiles showed clear peaks for all parameters, indicating that the inference algorithm performed well (Figure 3). Current and ancestral population size were estimated at *θ* = 42.24 (95% CI: 25.76 – 88.38) and *θ*_anc_ = 12.55 (95% CI: 5.25 – 21.41). Using a mutation rate of 4.23 × 10^−4^, these estimates correspond to *N*_e_ = 24,964 (95% CI: 15,224 – 52,234) and *N*_anc_ = 7,417 (95% CI: 3,103 – 12,653), respectively. *N*_ratio_ was estimated at 3.37 (95% CI: 1.55 – 7.10), indicating a significant population expansion. This demographic event appears quite recent as the timing since the beginning of this expansion was dated at Dg/2N = 0.0159 (95% CI: 0.0012 – 0.0830), which corresponds to *T* = 14,289 years ago (95% CI: 1,078 – 74,592), assuming a generation time of 9 years.

**Figure 3:**
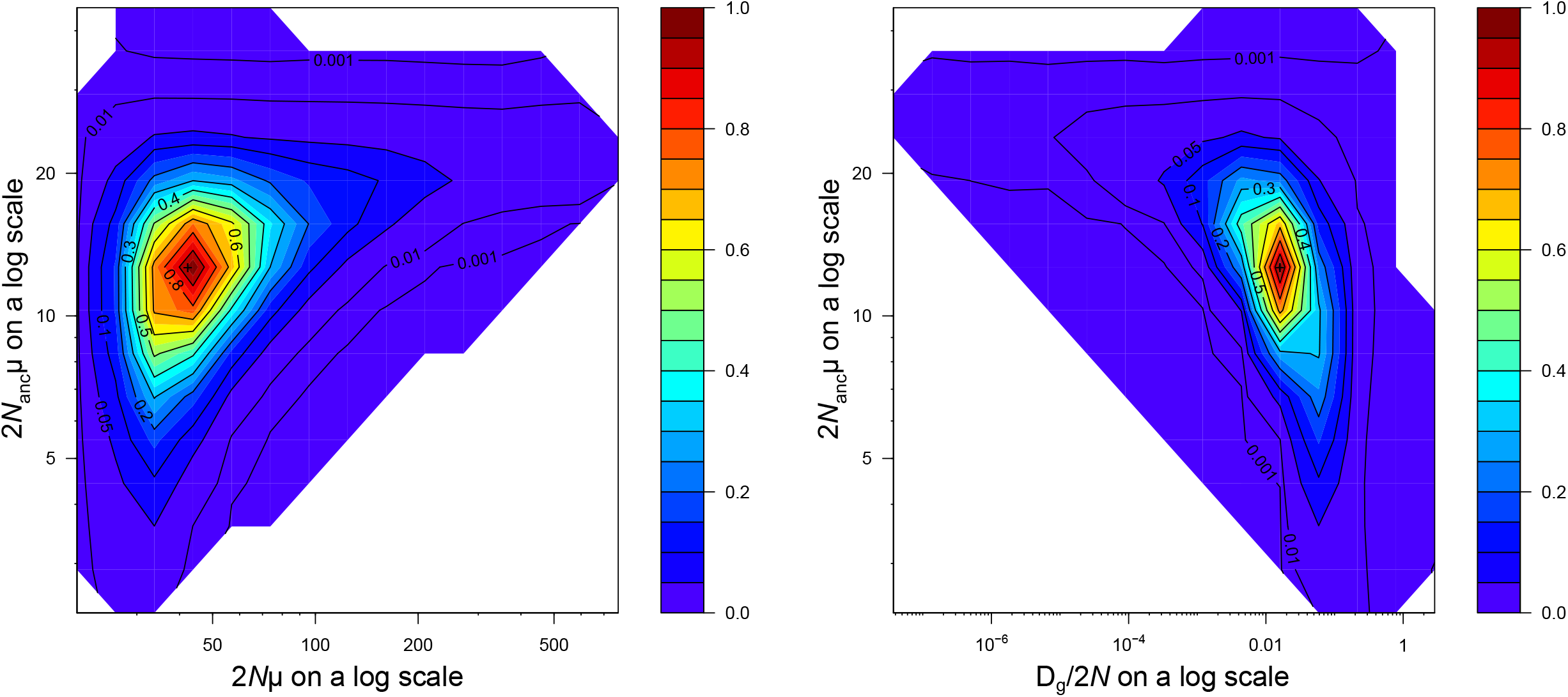
Pairwise likelihood-ratio profiles for South African *Carcharias taurus* inferred by MIGRAINE for *θ*_anc_ (2*N*_anc_μ): ancestral population size, *θ* (2*N*μ): current population size, *D* (*D*_g_*/*2*N*): timing of demographic event. The profiles show a significant demographic expansion in the population. Axes are on a logarithmic scale. See Results for point estimates and associated 95% confidence intervals.

## Discussion

Estimates of contemporary effective population size may assist in guiding management decisions for exploited and vulnerable elasmobranch populations, as they provide information on the size of the breeding population and its genetic health^18,24^. Here, two methods used to reconstruct the demographic history of *Carcharias taurus* in South Africa provided no evidence for recent, anthropogenic population decline, and instead, they both identified a historical population expansion with parameter estimates that were largely congruent between the two methods. The scenario recovered here may undoubtedly oversimplify the true demographic history of this population but, as advocated by Cabrera and Palsbøll^54^, it is reasonable to focus on a few simple, contrasting demographic scenarios that capture key demographic events. Based on the ABC analyses, the population expansion occurred ~32,960 years ago, where a historic *N*_e_ of only 1,100 individuals expanded to a current effective size of approximately 15,900 individuals. Results obtained through IS are concordant with a strong demographic expansion and inferred higher population size estimates (*N*_e_ = 20,978 and *N*_anc_ = 6,842), but with 95% CIs that broadly overlapped with those of the ABC results. The timing of this event was dated more recently at ~14,274 years ago, but again, 95% CIs from both methods overlapped broadly, ranging from ~2,097 to 87,750 (ABC) and ~1,722 – 71,669 (IS) years. Considering the wide confidence ranges and the uncertainty concerning some parameters (e.g. mutation rate, generation time), it is not possible to clearly link this demographic expansion to any specific geological or climatic events, but it is likely a result of climate oscillations during the last glacial cycle of the late Pleistocene^55^. Cycles of global climate change have caused widespread changes in the distribution and abundance of flora and fauna, leaving distinct genetic signatures of glacial refugia and post-colonization events in contemporary populations^56–58^. Despite earlier assumptions that highly mobile marine species may have been less affected by fluctuating climate conditions, expansions of elasmobranch populations following the last glacial maximum (~26,500 – 19,000 years ago) are commonly reported^59–62^. *Carcharias taurus* is a coastal species, and migratory movements are restricted by limits to its thermal tolerance^63,64^. Hence, it is likely that severe changes in oceanographic dynamics during climate oscillations have led to demographic changes and re-distributions of *C. taurus* in the southern hemisphere. Although we cannot link the expansion observed here to a specific climatic phase, it is clear that this event is unrelated to human activities.

Estimates of *N*_e_ and the absence of evidence for a recent, anthropogenic decline support the notion that the South African population is relatively healthy, as previously indicated by mark-recapture and beach meshing catch data^34,36^. Estimates of *N*_e_ found here closely approximate the mean estimate of the number of adult sharks obtained by mark-recapture data (16,700)^36^. This is in agreement with previous findings of high *N*_e_/*N*_c_ ratios in the East Australian *C. taurus* population^65,66^ and other elasmobranchs, such as the zebra shark, *Stegostoma fasciatum*^17^ and the sandbar shark, *Carcharhinus plumbeus*^18^. The observed pattern can be explained by similar life history traits, especially age at maturity and adult lifespan, which were found to account for half of the variation found in *N*_e_/*N*_c_ ratios across 63 animal and plant species^19^. Moreover, theoretical and empirical evidence show that high variability in reproductive success can strongly reduce *N*_e_/*N*_c_ ratios^67–69^. Taking into account that the reproductive output in *C. taurus* is small and constant (i.e. a maximum of two pups per litter as a result of intrauterine cannibalism)^70^, the reproductive variance is expected to be very small and this should increase the *N*_e_/*N*_c_ ratio compared to organisms with higher fecundity, such as teleost fish.

Exactly what size of *N*_e_ is needed to retain evolutionary potential is still controversial. Guidelines suggest that an *N*_e_ of at least 50 is required to prevent inbreeding depression^71^, and an *N*_e_ greater than 1,000 is needed to avoid the accumulation of deleterious alleles^72^. In order to retain the long-term evolutionary potential of a species, it has been suggested that an *N*_e_ greater than 500 or even 1,000 is necessary^73–75^. Although estimates of *N*_e_ were derived with relatively large 95% CIs, it can be concluded that the South African *C. taurus* population is nowhere near the beginning of genetic erosion. This is in stark contrast to the East Australian population, where a recent study found *N*_e_ to be as small as ~400 individuals^25^. The species gained a bad reputation in Australia due to its fierce appearance and was culled by spear and line fishers, resulting in severe population declines^76^. Despite recent conservation efforts, the Australian *C. taurus* populations are still under threat from incidental catch by recreational and commercial fishers, as well as beach meshing^76^. These threats, in combination with the species’ extremely low reproductive output, seem to have prevented a recovery, particularly along the East coast. Industrial fisheries started in South Africa around 1890, but the exploitation of marine resources commenced as long ago as the arrival of Dutch settlers in the mid-seventeenth century^77^. *Carcharias taurus* is susceptible to being caught as bycatch by small-scale coastal fisheries and the beach meshing program due its preference for shallow waters, but the species was never specifically targeted, and decommercialized through the Marine Living Resources Act in 1998^78^. This could explain the lack of evidence for a more recent population decline and indicates that anthropogenic pressure was not high enough to significantly reduce the size of this population. Given the critical status of this shark across much of its distribution, the South African *C. taurus* population may be the last remaining healthy population globally.

## Data Availability Statement

The multilocus genotype data file generated and analysed in this study is available in the Supplementary Information online.

## Acknowledgements

Special thanks are given to all volunteer anglers who collected samples as part of the Port Elizabeth Museum (PEM) co-operative shark tagging programme, as well as the PEM Director and staff for their support and infrastructure. Raphaël Leblois gave valuable advice on the use of the program MIGRAINE. This work is based on research supported in part by the National Research Foundation of South Africa through a Thuthuka grant to K.L.M. (Unique Grant no. 99440) and the University of Johannesburg (FRC/URC Grant to P.R.T.). J.D.K. is grateful to the University of Johannesburg for awarding her a Global Excellence and Stature (GES) PhD bursary.

## Author contributions

K.L.M. and M.L.D. contributed to the conception of the study and provided tissue samples. J.D.K. performed laboratory work and statistical analyses with guidance from P.R.T., A.E.B. and A.E. J.D.K. drafted the manuscript with assistance from P.R.T. and all authors revised previous versions of the manuscript. All authors approved the submitted version of the manuscript.

## References

1. Gabriel, W. & Bürger, R. Survival of small populations under demographic stochasticity. Theor. Popul. Biol. 41, 44–71 (1992).

2. Lande, R. Risks of population extinction from demographic and environmental stochasticity and random catastrophes. Am. Nat. 142, 911–927 (1993).

3. Melbourne, B. A. & Hastings, A. Extinction risk depends strongly on factors contributing to stochasticity. Nature 454, 100–103 (2008).

4. Frankham, R. Genetics and extinction. Biol. Conserv. 126, 131–140 (2005).

5. Spielman, D., Brook, B. W. & Frankham, R. Most species are not driven to extinction before genetic factors impact them. Proc. Natl. Acad. Sci. USA 101, 15261–15264 (2004).

6. Wright, S. Evolution in mendelian populations. Genetics 16, 97 (1931).

7. Nunney, L. The influence of mating system and overlapping generations on effective population size. Evolution 47, 1329–1341 (1993).

8. Ardren, W. R. & Kapuscinski, A. R. Demographic and genetic estimates of effective population size (*N*_e_) reveals genetic compensation in steelhead trout. Mol. Ecol. 12, 35–49 (2003).

9. Turner, T. F., Osborne, M. J., Moyer, G. R., Benavides, M. A. & Alò, D. Life history and environmental variation interact to determine effective population to census size ratio. Proc. R. Soc. B Biol. Sci. 273, 3065–3073 (2006).

10. Luikart, G., Ryman, N., Tallmon, D. A., Schwartz, M. K. & Allendorf, F. W. Estimation of census and effective population sizes: the increasing usefulness of DNA-based approaches. Conserv. Gen. 11, 355–373 (2010).

11. Palstra, F. P. & Fraser, D. J. Effective/census population size ratio estimation: a compendium and appraisal. Ecol. Evol. 2, 2357–2365 (2012).

12. Palstra, F. P. & Ruzzante, D. E. Genetic estimates of contemporary effective population size: what can they tell us about the importance of genetic stochasticity for wild population persistence? Mol. Ecol. 17, 3428–3447 (2008).

13. Nunney, L. Measuring the ratio of effective population size to adult numbers using genetic and ecological data. Evolution 49, 389–392 (1995).

14. Hauser, L. & Carvalho, G. R. Paradigm shifts in marine fisheries genetics: ugly hypotheses slain by beautiful facts. Fish Fish. 9, 333–362 (2008).

15. Ruggeri, P. et al. Coupling demographic and genetic variability from archived collections of European anchovy (*Engraulis encrasicolus*). Plos One 11, e0151507 (2016).

16. Waples, R. S., Grewe, P. M., Bravington, M. W., Hillary, R. & Feutry, P. Robust estimates of a high *N*_e_/*N* ratio in a top marine predator, southern bluefin tuna. Sci. Adv. 4, eaar7759 (2018).

17. Dudgeon, C. L. & Ovenden, J. R. The relationship between abundance and genetic effective population size in elasmobranchs: an example from the globally threatened zebra shark *Stegostoma fasciatum* within its protected range. Conserv. Gen. 16, 1443–1454 (2015).

18. Portnoy, D. S., McDowell, J. R., McCandless, C. T., Musick, J. A. & Graves, J. E. Effective size closely approximates the census size in the heavily exploited western Atlantic population of the sandbar shark, *Carcharhinus plumbeus*. Conserv. Gen. 10, 1697–1705 (2009).

19. Waples, R. S., Luikart, G., Faulkner, J. R. & Tallmon, D. A. Simple life-history traits explain key effective population size ratios across diverse taxa. Proc. R. Soc. B Biol. Sci. 280, 20131339 (2013).

20. Chevolot, M., Hoarau, G., Rijnsdorp, A. D., Stam, W. T. & Olsen, J. L. Phylogeography and population structure of thornback rays (*Raja clavata* L., Rajidae). Mol. Ecol. 15, 3693–3705 (2006).

21. Nunney, L. & Elam, D. R. Estimating the effective population size of conserved populations. Conserv. Biol. 8, 175–184 (1994).

22. Grimm, A., Gruber, B., Hoehn, M., Enders, K. & Henle, K. A model-derived short-term estimation method of effective size for small populations with overlapping generations. Methods Ecol. Evol. 7, 734–743 (2016).

23. Blower, D. C., Riginos, C. & Ovenden, J. R. NEOGEN: a tool to predict genetic effective population size (*N*_e_) for species with generational overlap and to assist empirical *N*_e_ study design. Mol. Ecol. Resour. 19, 260–271 (2019).

24. Pazmiño, D. A., Maes, G. E., Simpfendorfer, C. A., Salinas-de-León, P. & van Herwerden, L. Genome-wide SNPs reveal low effective population size within confined management units of the highly vagile Galapagos shark (*Carcharhinus galapagensis*). Conserv. Genet. 18, 1151–1163 (2017).

25. Reid-Anderson, S., Bilgmann, K. & Stow, A. Effective population size of the critically endangered east Australian grey nurse shark *Carcharias taurus*. Mar. Ecol. Prog. Ser. 610, 137–148 (2019).

26. Vignaud, T. M. et al. Genetic structure of populations of whale sharks among ocean basins and evidence for their historic rise and recent decline. Mol. Ecol. 23, 2590–2601 (2014).

27. Bansemer, C. S. & Bennett, M. B. Retained fishing gear and associated injuries in the east Australian grey nurse sharks (*Carcharias taurus*): implications for population recovery. Mar. Freshwater Res. 61, 97–103 (2010).

28. Irigoyen, A. & Trobbiani, G. Depletion of trophy large-sized sharks populations of the Argentinean coast, south-western Atlantic: insights from fishers’ knowledge. Neotrop. Ichthyol. 14, (2016).

29. Chiaramonte, G., Domingo, A. & Soto, J. *Carcharias taurus* (Southwest Atlantic subpopulation). The IUCN Red List of Threatened Species. e.T63163A12625032 (2007).

30. Pollard, D., Gordon, I., Williams, S., Flaherty, A. & McAuley, R. *Carcharias taurus* (East coast of Australia subpopulation). The IUCN Red List of Threatened Species. e.T44070A10854830 (2003).

31. Walls, A. & Soldo, A. *Carcharias taurus* (Mediterranean subpopulation). The IUCN Red List of Threatened Species: e.T3854A48947509 (2015).

32. Green, M., Ganassin, C. & Reid, D. Report into the NSW Shark Meshing (Bather Protection) Program. New South Wales Department of Primary Industries (2009).

33. Reid, D. D., Robbins, W. D., & Peddemors, V. M. Decadal trends in shark catches and effort from the New South Wales, Australia, Shark Meshing Program 1950–2010. Mar. Freshwater Res. 62, 676–693 (2011).

34. Dudley, S. F. J. & Simpfendorfer, C. A. Population status of 14 shark species caught in the protective gillnets off KwaZulu-Natal beaches, South Africa, 1978 – 2003. Mar. Freshwater Res. 57, 225 (2006).

35. Dicken, M. L., Smale, M. J. & Booth, A. J. Long-term catch and effort trends in Eastern Cape Angling Week competitions. Afr. J. Mar. Sci. 34, 259–268 (2012).

36. Dicken, M., Booth, A. J. & Smale, M. J. Estimates of juvenile and adult raggedtooth shark (*Carcharias taurus*) abundance along the east coast of South Africa. Can. J. Fish. Aquat. Sci. 65, 621–632 (2008).

37. Klein, J. D., Bester-van der Merwe, A. E., Dicken, M. L., Mmonwa, K. L. & Teske, P. R. Reproductive philopatry in a coastal shark drives age-related population structure. Mar. Biol. 166, 26 (2019).

38. Van Oosterhout, C., Hutchinson, W. F., Wills, D. P. M. & Shipley, P. MICRO-CHECKER: software for identifying and correcting genotyping errors in microsatellite data. Mol. Ecol. Notes 4, 535–538 (2004).

39. Rousset, F. GENEPOP’007: a complete re-implementation of the GENEPOP software for Windows and Linux. Mol. Ecol. Res. 8, 103–106 (2008).

40. Benjamini, Y. & Yekutieli, D. The control of the false discovery rate in multiple testing under dependency. Ann. Stat. 29, 1165–1188 (2001).

41. R Core Team. R: a language and environment for statistical computing. R Foundation for Statistical Computing (2017).

42. Chikhi, L., Sousa, V. C., Luisi, P., Goossens, B. & Beaumont, M. A. The confounding effects of population structure, genetic diversity and the sampling scheme on the detection and quantification of population size changes. Genetics 186, 983–995 (2010).

43. Leblois, R. et al. Maximum-likelihood inference of population size contractions from microsatellite data. Mol. Biol. Evol. 31, 2805–2823 (2014).

44. Peakall, R. & Smouse, P. E. GenAlEx 6.5: genetic analysis in Excel. Population genetic software for teaching and research-an update. Bioinformatics 28, 2537–2539 (2012).

45. Cornuet, J. M. et al. DIYABC v2.0: A software to make approximate Bayesian computation inferences about population history using single nucleotide polymorphism, DNA sequence and microsatellite data. Bioinformatics 30, 1187–1189 (2014).

46. Goldman, K. J., Branstetter, S. & Musick, J. A. A re-examination of the age and growth of sand tiger sharks, *Carcharias taurus*, in the western North Atlantic: the importance of ageing protocols and use of multiple back-calculation techniques. Environ. Biol. Fish 77, 241–252 (2006).

47. Garza, J. C. & Williamson, E. G. Detection of reduction in population size using data from microsatellite loci. Mol. Ecol. 10, 305–318 (2001).

48. Cornuet, J.-M. et al. Inferring population history with DIY ABC: a user-friendly approach to approximate Bayesian computation. Bioinformatics 24, 2713–2719 (2008).

49. de Irio, M. D. & Griffiths, R. C. Importance sampling on coalescent histories. I. Adv. Appl. Prob. 36, 417–433 (2004).

50. de Iorio, M. D. & Griffiths, R. C. Importance sampling on coalescent histories. II: subdivided population models. Adv. Appl. Prob. 36, 434–454 (2004).

51. Cornuet, J. M. & Beaumont, M. A. A note on the accuracy of PAC-likelihood inference with microsatellite data. Theor. Popul. Biol. 71, 12–19 (2007).

52. Nance, H. A., Klimley, P., Galván-Magaña, F., Martínez-Ortíz, J. & Marko, P. B. Demographic processes underlying subtle patterns of population structure in the scalloped hammerhead shark, *Sphyrna lewini*. Plos One 6, e21459 (2011).

53. Vignaud, T. M. et al. Blacktip reef sharks, *Carcharhinus melanopterus*, have high genetic structure and varying demographic histories in their Indo-Pacific range. Mol. Ecol. 23, 5193–5207 (2014).

54. Cabrera, A. A. & Palsbøll, P. J. Inferring past demographic changes from contemporary genetic data: A simulation-based evaluation of the ABC methods implemented in DIYABC. Mol. Ecol. Res. 17, e94–e110 (2017).

55. Blunier, T. & Brook, E. J. Timing of millennial-scale climate change in Antarctica and Greenland during the last glacial period. Science 291, 109–112 (2001).

56. Fraser, C. I., Nikula, R., Ruzzante, D. E. & Waters, J. M. Poleward bound: biological impacts of southern hemisphere glaciation. Trends Ecol. Evol. 27, 462–471 (2012).

57. Provan, J. & Bennett, K. Phylogeographic insights into cryptic glacial refugia. Trends in Ecol. Evol. 23, 564–571 (2008).

58. Tolley, K., Groeneveld, J., Gopal, K. & Matthee, C. Mitochondrial DNA panmixia in spiny lobster *Palinurus gilchristi* suggests a population expansion. Mar. Ecol. Prog. Ser. 297, 225–231 (2005).

59. O’Brien, S. M., Gallucci, V. F. & Hauser, L. Effects of species biology on the historical demography of sharks and their implications for likely consequences of contemporary climate change. Conserv. Gen. 14, 125–144 (2013).

60. Portnoy, D. S. et al. Contemporary population structure and post-glacial genetic demography in a migratory marine species, the blacknose shark, *Carcharhinus acronotus*. Mol. Ecol. 23, 5480–5495 (2014).

61. Domingues, R. R., Hilsdorf, A. W. S., Shivji, M. M., Hazin, F. V. H. & Gadig, O. B. F. Effects of the Pleistocene on the mitochondrial population genetic structure and demographic history of the silky shark (*Carcharhinus falciformis*) in the western Atlantic Ocean. Rev. Fish Biol. Fish. 28, 213–227 (2018).

62. Portnoy, D. S. et al. Population structure, gene flow, and historical demography of a small coastal shark (*Carcharhinus isodon*) in US waters of the Western Atlantic Ocean. ICES J. Mar. Sci. 73, 2322–2332 (2016).

63. Smale, M. J., Booth, A. J., Farquhar, M. R., Meÿer, M. R. & Rochat, L. Migration and habitat use of formerly captive and wild raggedtooth sharks (*Carcharias taurus*) on the southeast coast of South Africa. Mar. Biol. Res. 8, 115–128 (2012).

64. Bradshaw, C. J. A., Peddemors, V. M., McAuley, R. B. & Harcourt, R. G. Population viability of eastern Australian grey nurse sharks under fishing mitigation and climate change. Final Report to the Commonwealth of Australia, Department of the Environment, Water, Heritage and the Arts (2008).

65. Ahonen, H., and Stow, A. Population size and structure of grey nurse shark in east and west Australia. Final Report to Department of Marine and Freshwater Environment, Water, Heritage and the Arts (2009).

66. Otway, N. M. & Burke, A. L. Mark-recapture population estimate and movements of grey nurse sharks. NSW Fisheries Final Report Series No. 63 (2004).

67. Belmar-Lucero, S. et al. Concurrent habitat and life history influences on effective/census population size ratios in stream-dwelling trout. Ecol. Evol. 2, 562–573 (2012).

68. Frankham, R. Effective population size/adult population size ratios in wildlife: a review. Genet. Res. 66, 95 (1995).

69. Hedgecock, D. Does variance in reproductive success limit effective population sizes of marine organisms in Genetics and Evolution of Aquatic Organisms (ed. Beaumont, A.) 122–134 (Springer, 1994).

70. Gilmore, R., Putz, O. & Dodrill, J. Oophagy, intrauterine cannibalism and reproductive strategy in lamnoid sharks in Reproductive Biology and Phylogeny of Chondrichthyes (ed. Hamlett, W.) 435–463 (Science Enfield, NH: Publishers Inc., 2005).

71. Franklin, I. R. Evolutionary change in small populations in Conservation Biology: an Evolutionary-Ecological Perspective 135–150 (Sinauer, 1980).

72. Palstra, F. P. & Ruzzante, D. E. Genetic estimates of contemporary effective population size: what can they tell us about the importance of genetic stochasticity for wild population persistence? Mol. Ecol. 17, 3428–3447 (2008).

73. Frankham, R. Challenges and opportunities of genetic approaches to biological conservation. Biol. Conserv. 143, 1919–1927 (2010).

74. Franklin, I. R. & Frankham, R. How large must populations be to retain evolutionary potential? Anim. Conserv. 1, 69–70 (1998).

75. Frankham, R., Bradshaw, C. J. A. & Brook, B. W. Genetics in conservation management: revised recommendations for the 50/500 rules, Red List criteria and population viability analyses. Biol. Conserv. 170, 56–63 (2014).

76. Department of the Environment. Recovery plan for the grey nurse shark (Carcharias taurus) in Australia (2014).

77. van Sittert, L. The marine fisheries of South Africa in Oxford Research Encyclopedia of African History 1–15 (2017).

78. RSA (Republic of South Africa). Marine living resources act (act no. 18 of 1998). 1–67 (1998).

