## Supplementary Information for "A globally threatened shark, *Carcharias taurus*, shows no population decline in South Africa"

### Analysis of population structure

**Table S1:** Population pairwise comparison based on 12 microsatellite markers and seven sampling sites for *Carcharias taurus* (N = 189). *F*_ST_ values are below the diagonal, *D*_est_ values are above the diagonal. Values in bold indicate significance before applying B-Y method for multiple comparison.

|  | SC | JB | PE | PA | EL | ZIN | RB |
| --- | --- | --- | --- | --- | --- | --- | --- |
| SC | -- | -0.027 | 0.004 | -0.001 | -0.020 | 0.012 | -0.002 |
| JB | 0.017 | -- | -0.029 | **0.048** | 0.013 | -0.014 | 0.001 |
| PE | 0.021 | 0.012 | -- | 0.028 | 0.004 | 0.003 | 0.018 |
| PA | 0.021 | **0.020** | 0.020 | -- | 0.033 | **0.046** | **0.044** |
| EL | 0.014 | 0.011 | 0.012 | 0.014 | -- | 0.019 | -0.002 |
| ZIN | 0.018 | 0.009 | 0.013 | **0.017** | 0.008 | -- | 0.005 |
| RB | 0.015 | 0.009 | 0.013 | **0.015** | 0.005 | 0.007 | -- |

SC = Sedgefield, JB = Jeffreys Bay, PE = Port Elizabeth, PA = Port Alfred, ZIN = Zinkwazi, RB = Richards Bay

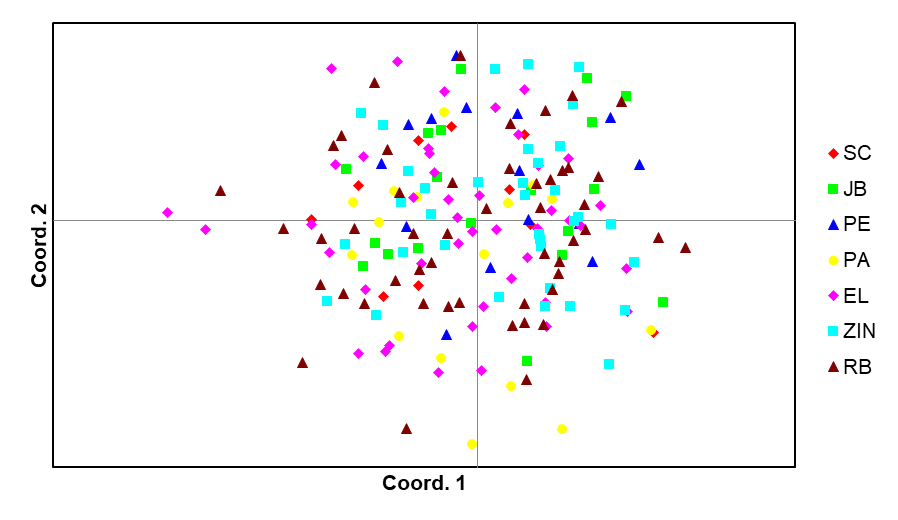

**Figure S1:** Principal Coordinate Analysis (PCoA) based on 189 samples of *Carcharias taurus* collected at seven sites along the South African coast and genotyped for 12 microsatellite markers*.* SC = Sedgefield, JB = Jeffreys Bay, PE = Port Elizabeth, PA = Port Alfred, ZIN = Zinkwazi, RB = Richards Bay

### DIYABC: preliminary analysis and determination of suitable priors

The prior distribution for contemporary *N*_e_ in the initial ABC analysis was based on the estimate of abundance reported by Dicken et al.^1^. Microsatellite loci were assumed to follow a generalized stepwise mutation model (GSM)^2^ as it is considered the most suitable mutation model for microsatellites and reduces the risk of false positives in bottleneck testing^3,4^. Under the GSM, the mean mutation rate (mean µ) and the mean parameter of the geometric distribution (mean *P*) of length in terms of the number of repeats of mutation events were estimated. Rates for microsatellite mutation rates used in the literature vary widely, and may range from 1.0 × 10^-2^ to 9.0 × 10^-6^ per locus per generation^5,6^. Molecular evolution is slower in elasmobranch than in mammals^7,8^, hence previous studies have used mutation rates at the lower end of this range due to the lack of elasmobranch-specific rates^9,10^. We accounted for this uncertainty by selecting a wide prior for µ at individual loci. Prior distributions for all parameters are shown in Table S2.

**Table S2:** Demographic and microsatellite mutation model prior distributions and resulting posterior estimates of estimate effective population size and microsatellite mutation rate of initial approximate Bayesian computation simulation one panmictic population.

| **Parameter** | **Prior distribution [min., max.]** | **Posterior distribution**  **[95% HPDI]** |
| --- | --- | --- |
| N1 | UN [10^1^ – 3 × 10^4^] | 13,700 [4.74 × 10^3^ – 2.88 × 10^4^] |
| µ | UN [1 × 10^-5^ – 1 × 10^-3^] | 4.23 × 10^-4^ [1.37 × 10^-4^ – 9.61 × 10^-4^] |
| µ_ind_ | GA [1 × 10^-6^ – 1 × 10^-2^, mean: 1 × 10^-4^] | - |
| P | UN [1 × 10^-1^ – 1] | 4.27 × 10^-1^ [1.56 × 10^-1^ – 8.99 × 10^-1^] |
| P_ind_ | GA [1 × 10^-1^ – 1, mean: 2.2 × 10^-1^] | - |

N1 = present effective population size, µ = microsatellite mutation rate over all loci, µ_ind_ = microsatellite mutation rate at individual loci, P = parameter of the geometric distribution over all loci, P_ind_ = parameter of the geometric distribution at individual loci, UN = uniform distribution, GA = gamma distribution, HPDI = highest posterior density interval.

A principal component analysis (PCA) was computed using 10,000 simulated data sets with the observed data added on each plane. This first assessment showed that the selected prior distributions produced simulated data that are close enough to the observed data (Figure S2), therefore median estimates of demographic and genetic parameters were used as a basis for the prior distributions in the following comparison of different possible demographic models (Table S3).

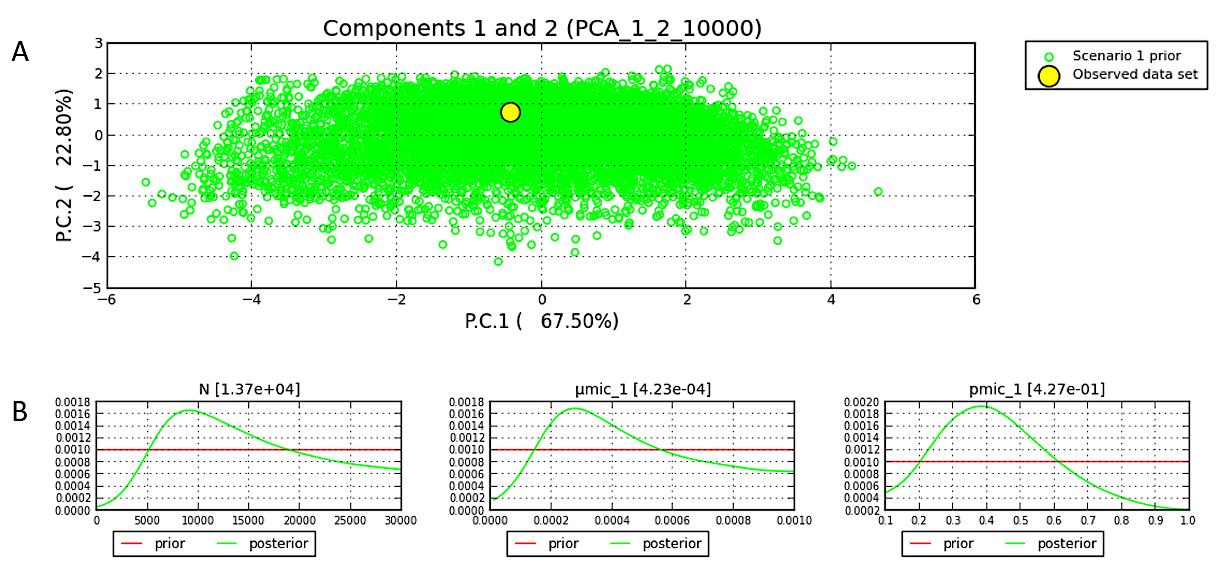

**Figure S2**: Principal component analysis in the space of 10,000 datasets simulating a panmictic, stable population (open circles) with the observed data added on each plane (yellow circle) (A) and posterior distributions of demographic and genetic parameters (B).

**Table S3:** Demographic and microsatellite mutation model prior distributions for approximate Bayesian computation model comparison.

|  | **Analysis 1** | **Analysis 2** |
| --- | --- | --- |
| **Parameter** | **Prior distribution [min, max]** | |
| N1 | N [8 × 10^3^ – 2.5 × 10^4^, mean: 1.37 × 10^4^, sd: 3 ×10^3^] | N [8 × 10^3^ – 2.5 × 10^4^, mean: 1.37× 10^4^, sd: 3 × 10^3^] |
| *t* | **UN [****1 – 5** × **10^3^]** | **UN [1 – 50]** |
| N2 | UN [10^2^ – 7 × 10^3^] | UN [10^2^ – 7 × 10^3^] |
| N3 | UN [3.2 × 10^4^ – 2 × 10^5^] | UN [3.2 × 10^4^ – 2 × 10^5^] |
| µ | GA [1.37 × 10^-4^ – 9.61 × 10^-4^, mean: 4.23 × 10^-4^] | GA [1.37 × 10^-4^ – 9.61 × 10^-4^, mean: 4.23 × 10^-4^] |
| *P* | GA [1.56 × 10^-1^ – 8.99 × 10^-1^, mean: 4.27 × 10^-1^] | GA [1.56 × 10^-1^ – 8.99 × 10^-1^, mean: 4.27 × 10^-1^] |

N1 = current effective population size, *t* = time of population size change in generations, N2 = reduced historical effective population size, N3 = large historical effective population size, µ = microsatellite mutation rate (over all loci and individual loci), *P* = parameter of the geometric distribution (over all loci and individual loci), sd = standard deviation for normally distributed priors, N = normal distribution, UN = uniform distribution, GA = gamma distribution. See Figure 1 for a visualization of the three scenarios. Note that analysis 1 and 2 differed only in the restriction of the parameter *t*.

### DIYABC: model evaluation

A first pre-evaluation through PCA showed that prior distributions produced simulated data close to the observed data (Figure S3). For analysis 1, which included a wide prior for the time at which the size of the population changed (*t*), scenario 2 (population expansion), had the highest posterior probability which was close to the maximum possible value of 1.0 with very narrow 95% confidence intervals (Figure 2a). When *t* was restricted to the past ~400 years (analysis 2), scenario 1 (constant population size) had the highest support when estimating posterior probabilities (Figure 2b). In both analyses, scenario 3 (population decline) received no support. Because simulations with *t* restricted to the last ~400 years did not provide conclusive posterior distributions as much of the genetic signal present in the data was excluded, estimates of demographic parameters were derived from scenario 2 simulated in analysis 1. Precision of the estimates was evaluated by simulating 1,000 datasets based on a fixed parameter set (‘true values’). The parameters of these pseudo-observed datasets (pods) were estimated using the previously 10^6^ simulations based on the prior information. The estimates were then compared to the true values used to generate the test data sets. This allowed to assess the quality of the point estimators by computing the mean relative bias (MRB). Precision of the posterior distributions was further evaluated by calculating the square root of the relative mean square error (RRMSE) and the proportion of test data sets for which the 95% credibility intervals included the true value (95% coverage). Small MRB and RRMSE values in Table S4 demonstrated that all posterior median estimates of parameters had small bias and high precision. Similarly, precision of the posterior distributions was high, as indicated by the high proportion of test data sets among which the true value was within the 95% credibility interval (Table S4).

In a next step, the “goodness-of-fit” of the different models and their parameter posterior distributions were assessed by generating 1,000 datasets, randomly drawing parameters from the posterior distributions. This posterior predictive distribution was then used in a PCA, together with datasets simulated from the prior distribution as well as the observed data added to each plane. Figure S4 shows that scenario 2 fits the observed data well, with the observed data surrounded by a small cluster of datasets from the posterior predictive distribution and a wider cloud of data sets simulated from the prior.

As a last measure to evaluate confidence in the model choice, simulated pods were used to estimate classification error (type I error) and misclassification error (type II error). The type I error was estimated as the proportion of pods among which the true model was incorrectly excluded, whereas type II error was estimated as the proportion of pods among which an incorrect model was recovered. The type I error (probability that scenario 2 is rejected although it is the true scenario) was 0.080 and 0.066 in the direct and logistic approach respectively. Type II error (probability of choosing scenario 2 when it is not the true scenario) was 0.038 and 0.056 in the direct and logistic approach respectively.
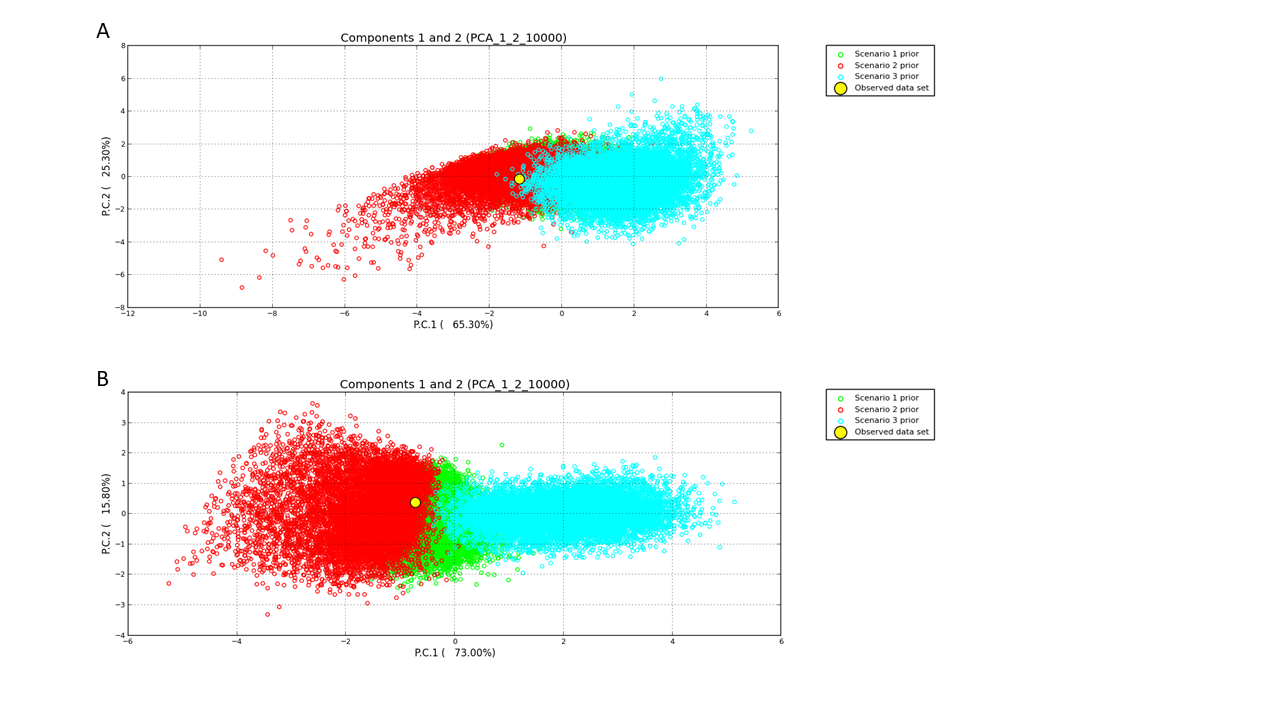

**Figure S3:** Principal component analysis in the space of 10,000 simulated data sets of each scenario (open circles) with the observed data added on each plane (yellow circle) for (A) analysis 1 and (B) analysis 2.

**Table S4:** Posterior estimates of demographic and genetic parameters estimated from scenario 2.

| Parameter | median | 95% HPDI | MRB | RRMSE | 95% cov. |
| --- | --- | --- | --- | --- | --- |
| N1 | 1.59 × 10^4^ | 9.81 × 10^3^ – 2.16 × 10^4^ | -0.065 | 0.089 | 1.000 |
| *t* | 4.12 × 10^3^ | 6.69 × 10^2^ – 9.42 × 10^3^ | 0.175 | 0.351 | 1.000 |
| N2 | 1.10 × 10^3^ | 1.43 × 10^2^ – 5.81 × 10^3^ | 1.703 | 1.934 | 1.000 |
| µ | 4.57 × 10^-4^ | 1.77 × 10^-4^ – 8.98 × 10^-4^ | -0.158 | 0.215 | 1.000 |
| *P* | 5.19 × 10^-1^ | 2.67 × 10^-1^ – 8.55 × 10^-1^ | -0.1011 | 0.233 | 0.984 |

N1 = present effective population size, *t* = time of population size change in generations, N2 = historical effective population size, µ = microsatellite mutation rate, *P* = parameter of the geometric distribution (over all loci and individual loci), HPDI = highest posterior density interval, MRB = mean relative bias, RRMSE = square root of the relative mean square error, 95% cov. = proportion of test data sets for which the 95% credibility intervals included the true value.

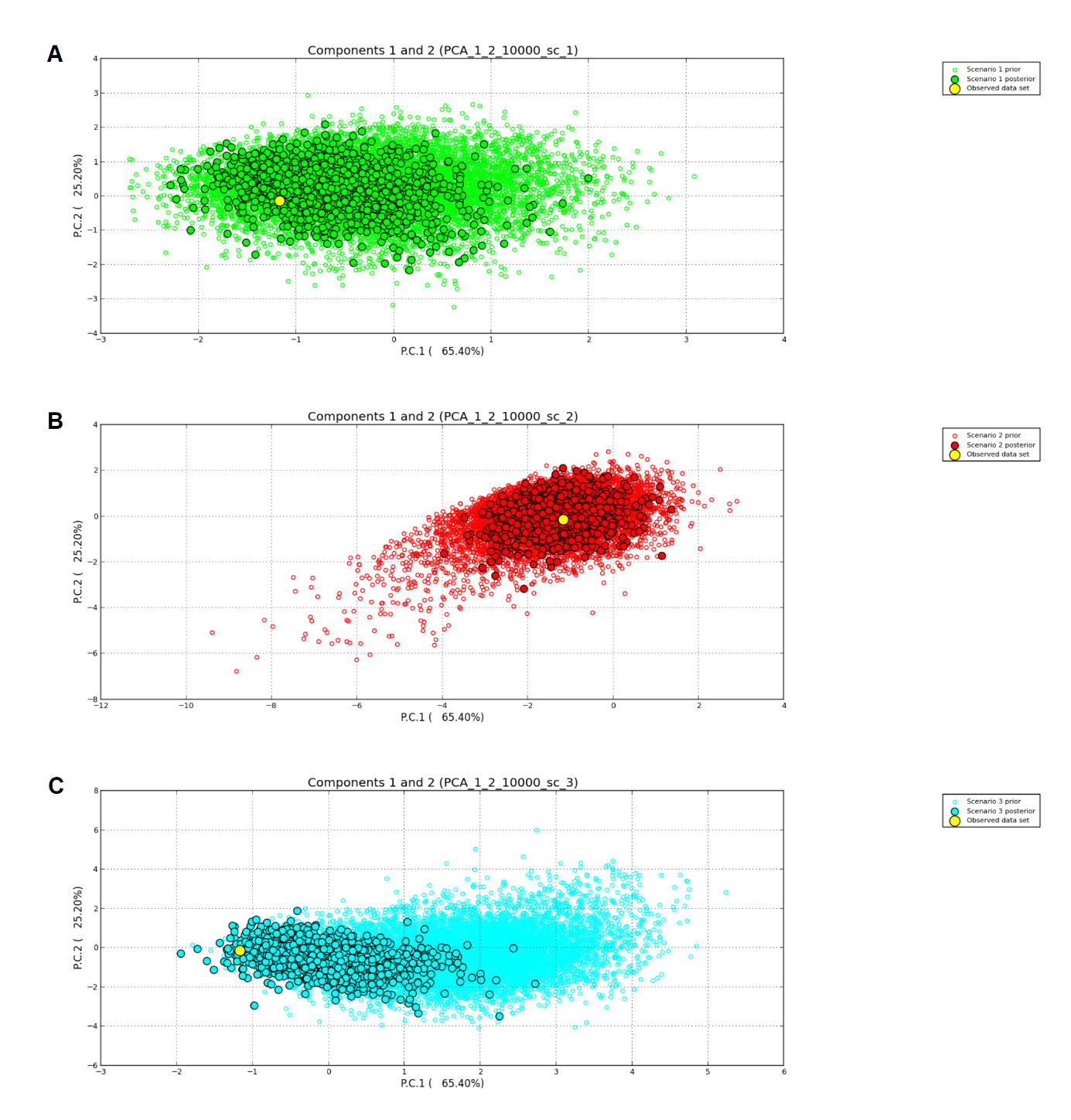

**Figure S4:** Evaluation of the different demographic models investigated within an approximate Bayesian framework. “Goodness-of-fit” and parameter posterior distributions were assessed by generating 1,000 datasets, randomly drawing parameters from the posterior distributions. This posterior predictive distribution (closed circles) is here used in a principal component analysis, together with datasets simulated from the prior distribution (open circles) as well as the observed data (yellow circle) added to each plane. A = stable population (scenario 1), B = population expansion after time step t (scenario 2), B = population decline after time step t (scenario 3).

### References

1.	Dicken, M. L., Booth, A. J. & Smale, M. J. Estimates of juvenile and adult raggedtooth shark (*Carcharias taurus*) abundance along the east coast of South Africa. *Can. J. Fish. Aquat. Sci.* **65**, 621–632 (2008).

2.	Estoup, A., Jarne, P. & Cornuet, J. M. Homoplasy and mutation model at microsatellite loci and their consequences for population genetics analysis. *Mol. Ecol.* **11**, 1591–1604 (2002).

3.	Leblois, R. *et al.* Maximum-likelihood inference of population size contractions from microsatellite data. *Mol. Biol. Evol.* **31**, 2805–2823 (2014).

4.	Peery, M. Z. *et al.* Reliability of genetic bottleneck tests for detecting recent population declines. *Mol. Ecol.* **21**, 3403–3418 (2012).

5.	Ellegren, H. Microsatellite mutations in the germline: implications for evolutionary inference. *Trends Genet.* **16**, 551–558 (2000).

6.	Zhivotovsky, L. A., Bennett, L., Bowcock, A. M. & Feldman, M. W. Human population expansion and microsatellite variation. *Mol. Biol. Evol.* **17**, 757–767 (2000).

7.	Martin, A. P., Naylor, G. J., & Palumbi, S. R. Rates of mitochondrial DNA evolution in sharks are slow compared with mammals. *Nature* **357**, 153 (1992).

8.	Martin, A. P. Substitution rates of organelle and nuclear genes in sharks: implicating metabolic rate (again). *Mol. Biol. Evol.* **16**, 996–1002 (1999).

9.	Karl, S. A., Castro, A. L. F., Lopez, J. A., Charvet, P. & Burgess, G. H. Phylogeography and conservation of the bull shark (*Carcharhinus leucas*) inferred from mitochondrial and microsatellite DNA. *Conserv. Gen.* **12**, 371–382 (2011).

10.	Blower, D. C., Pandolfi, J. M., Bruce, B. D., Gomez-Cabrera, M. D. C. & Ovenden, J. R. Population genetics of Australian white sharks reveals fine-scale spatial structure, transoceanic dispersal events and low effective population sizes. *Mar. Ecol. Prog. Ser.* **455**, 229–244 (2012).
